# Detection and editing of the updated plastid- and mitochondrial-encoded proteomes for *Arabidopsis* with PeptideAtlas

**DOI:** 10.1101/2023.07.10.548362

**Authors:** Klaas J. van Wijk, Stephane Bentolila, Tami Leppert, Qi Sun, Zhi Sun, Luis Mendoza, Margaret Li, Eric W. Deutsch

## Abstract

*Arabidopsis thaliana* Col-0 has plastid and mitochondrial genomes encoding for over one hundred proteins and several ORFs. Public databases (*e.g.* Araport11) have redundancy and discrepancies in gene identifiers for these organelle-encoded proteins. RNA editing results in changes to specific amino acid residues or creation of start and stop codons for many of these proteins, but the impact of such RNA editing at the protein level is largely unexplored due to the complexities of detection. This study first assembled the non-redundant set of identifiers, their correct protein sequences, and 452 predicted non-synonymous editing sites of which 56 are edited at lower frequency. Accumulation of edited and/or unedited proteoforms was then determined by searching ∼259 million raw MSMS spectra from ProteomeXchange as part of Arabidopsis PeptideAtlas (www.peptideatlas.org/builds/arabidopsis/). All mitochondrial proteins and all except three plastid-encoded proteins (NDHG/NDH6, PSBM, RPS16), but none of the ORFs, were identified; we suggest that all ORFs and RPS16 are pseudogenes. Detection frequencies for each edit site and type of edit (*e.g.* S to L/F) were determined at the protein level, cross-referenced against the metadata (*e.g.* tissue), and evaluated for technical challenges of detection.167 predicted edit sites were detected at the proteome level. Minor frequency sites were indeed also edited at low frequency at the protein level. However, except for sites RPL5-22 and CCB382-124, proteins only accumulate in edited form (>98 –100% edited) even if RNA editing levels are well below 100%. This study establishes that RNA editing for major editing sites is required for stable protein accumulation.

## INTRODUCTION

*Arabidopsis thaliana* (from here on Arabidopsis) and all other plants have plastid and mitochondrial genomes encoding for proteins, several ORFs, tRNAs and rRNAs. These organelle-encoded proteins are components of the protein complexes of the plastid and mitochondrial electron transport chains, components of the transcriptional and translations machineries, protein biogenesis factors, as well as several proteins involved in metabolism (Green, 2011; Zoschke and Bock, 2018; Moller et al., 2021). The precise function of just a few organelle-encoded proteins is still unknown (*e.g.* plastid YCF1/YCF214 and YCF2), whereas the ORFs might be pseudogenes (*i.e.* plastid-encoded YCF15 and mitochondrial ORFX/TATC, ORF114 and ORF240A). Most sources report around 88 protein-coding plastid genes and 31-33 protein-coding mitochondrial genes in Arabidopsis, based on initial sequenced genomes for plastids for ecotype Columbia (Col-0) (Sato et al., 1999) and mitochondria for ecotype C24 (Unseld et al., 1997). The Araport11 genome assembly (and also the prior assembly TAIR10) includes 122 mitochondrial-encoded protein identifiers (ATMG) rather than the expected 31-33 (Sloan et al., 2018). Furthermore, several plastid-(ATCG) or mitochondrial (ATMG) –encoded protein identifiers in Araport11/TAIR10 represent individual exons that are trans-spliced to make complete protein-coding transcripts (*i.e.* plastid-encoded RPS12 and mitochondrial-encoded NDH1,2,5), or are redundant because they represent duplicates copies of protein-coding genes located on the inverse repeats (IRs) of the plastid genome. Additionally, many of the organelle-encoded proteins have more than one protein name in the literature and various databases. The current study aims to first clarify this confusion and provides a consensus non-redundant set of plastid– and mitochondrial encoded proteins with their amino acid sequences for ecotype Col-0, including the protein identifiers (ATCG and ATMG). We suggest that this updated set will be incorporated in the forthcoming new Arabidopsis genome annotation for Col-0 (<u>tinyurl.com/Athalianav12</u>).

Several plastid– and most mitochondrial-encoded mRNAs undergo C to U mRNA editing which can affect the resulting protein sequence, *i.e*. non-synonymous edits (Takenaka et al., 2013; Germain et al., 2015; Fuchs et al., 2020; Small et al., 2020). In the case of NDHD/NDH4, mRNA editing results in introduction of a start codon (thus generating a translatable mRNA) and in several cases, mitochondrial gene editing introduces an extra stop codon (which would result in a truncated protein). It is generally believed that RNA editing events are required for protein function and/or protein stability (Small et al., 2023). The extent of quantitative editing for each site has been reported at the RNA level and several editing sites showed variability in the extent of editing (partial editing) depending on, growth or stress condition, developmental state and/or tissue type (Ruwe et al., 2013; Germain et al., 2015). The impact of RNA editing on proteins is best done by proteomics and tandem mass spectrometry (MSMS). However, very little systematic information about editing is available at the protein level, mostly because of technical challenges and low sequence coverage. In a typical proteome experiment, the isolated proteome is converted into peptides using enzymatic digestion with a protease (typically trypsin), followed by tandem MS analysis of the peptide mixture. The technical challenges to map editing sites at the protein level includes incomplete protein sequence coverage, in particular for regions that are very hydrophobic (transmembrane domains) or with many positive amino acids (Lys, Arg). An added challenge is to accommodate the protein search space to allow for detection of closely spaced editing sites in the same protein independently of each other.

In this study we determined the accumulation of unedited and edited isoforms of the plastid– and mitochondrial-encoded proteins, and possible partial editing due to biological conditions by searching ∼259 million MSMS spectra as part of the Arabidopsis PeptideAtlas project (van Wijk et al., 2021; van Wijk et al., 2023). To facilitate the protein search and optimize the protein search space, we reached out to members of the plant community and gathered literature annotations (e.g. (Germain et al., 2015; Lenz et al., 2018)) to obtain the most complete set of possible organelle-encoded proteins and their amino acid sequences, including their unedited and edited variants. Whereas we identified most organelle-encoded proteins, no MSMS support was found for several of the ORFs suggesting these are indeed pseudogenes. We report on the detection of edited sites and make recommendations for protein search space and search strategies for future proteomics studies that include the plastid– and mitochondrial-encoded proteomes. Finally, we also report on four important physiological post-translation modifications of the organelle-encoded proteins in PeptideAtlas build 2, namely N-terminal acetylation, lysine acetylation, and phosphorylation, as well as ubiquitination, based on raw MS data from submissions in ProteomeXchange that specifically enriched for these PTMs.

## RESULTS

### Assembly of the non-redundant plastid and mitochondrial encoded proteins, protein sequences and predicted editing sites

To arrive at a consensus set of plastid– and mitochondrial-encoded protein sequences and their predicted non-synonymous RNA editing sites, we collected information from a wide range of public resources. Furthermore, we reached out to the plastid– and mitochondrial research community to solicit input on plastid– and mitochondrial editing sites as well as significance of predicted organellar ORFs (see Acknowledgements).

Table 1 summarizes the plastid-encoded proteins with their gene ATCG identifier, protein name and function, and non-synonymous editing events. These identifiers and their unedited and (partially) edited amino acid sequence variants can be downloaded at Arabidopsis PeptideAtlas. There are 88 ATCG identifiers in TAIR (identical across Araport11 and TAIR10) due to various types of redundances (see below), but there are only 79 unique full length protein sequences. Within the new release of the Arabidopsis PeptideAtlas and in this study, we removed this redundancy (Table 1). For two proteins (MATK and NDHB/NDH2) there are conflicting database entries about the start of the protein sequence, *e.g.* compare the shorter forms of MATK in NP_050140 and P56784 with the extended forms in TAIR10/Araport11 ATCG00040 (see Arabidopsis PeptideAtlas). Similarly compare the longer form of NDHB/NDH2 in NP_051103 and P0CC32 with the shorter form in Araport11/TAIR10 ATCG00890 (see Arabidopsis PeptideAtlas). In case of MATK the shorter form (22 amino acids shorter) appears correct (N-terminus is MDKFQGYLEF) and not MCHFRTQENKDFTFSSNRISIQ) as also supported by lack of observed matched peptides to the extended N-terminus. In case of NDHB/NDH2 the longer form (123 amino acids longer) is correct (N-terminus is MIWHVQNENF and not MAITEFLLF) (Table 1). For RPS12 there are three ATCG identifiers each representing an exon – and we selected the identifier with the lowest number (ATCG00065) and associated the full-length protein sequence with this identifier. Six pairs of ATCG identifiers describe identical duplicates of proteins expressed from genes in the two inverted repeats (IRs) (RPL2, RPL23, RPS7, YCF15, YCF2 and NDHB). For each pair, we selected the identifier with the lowest number and excluded the others. Finally, ATCG01000 represents a truncated, non-functional YCF1 ORF (pers comm. R Bock) and this entry can also be removed for proteome analysis (and there are no peptides that map uniquely to the ATCG01000 sequence). In the case of 17 (unique) proteins, mRNA editing can result in one or more amino acid changes. In total, predicted editing would result in 31 amino acid changes and generation of one start methionine in case of NDHD/NDH4 (Table 1).

**Table 1.**
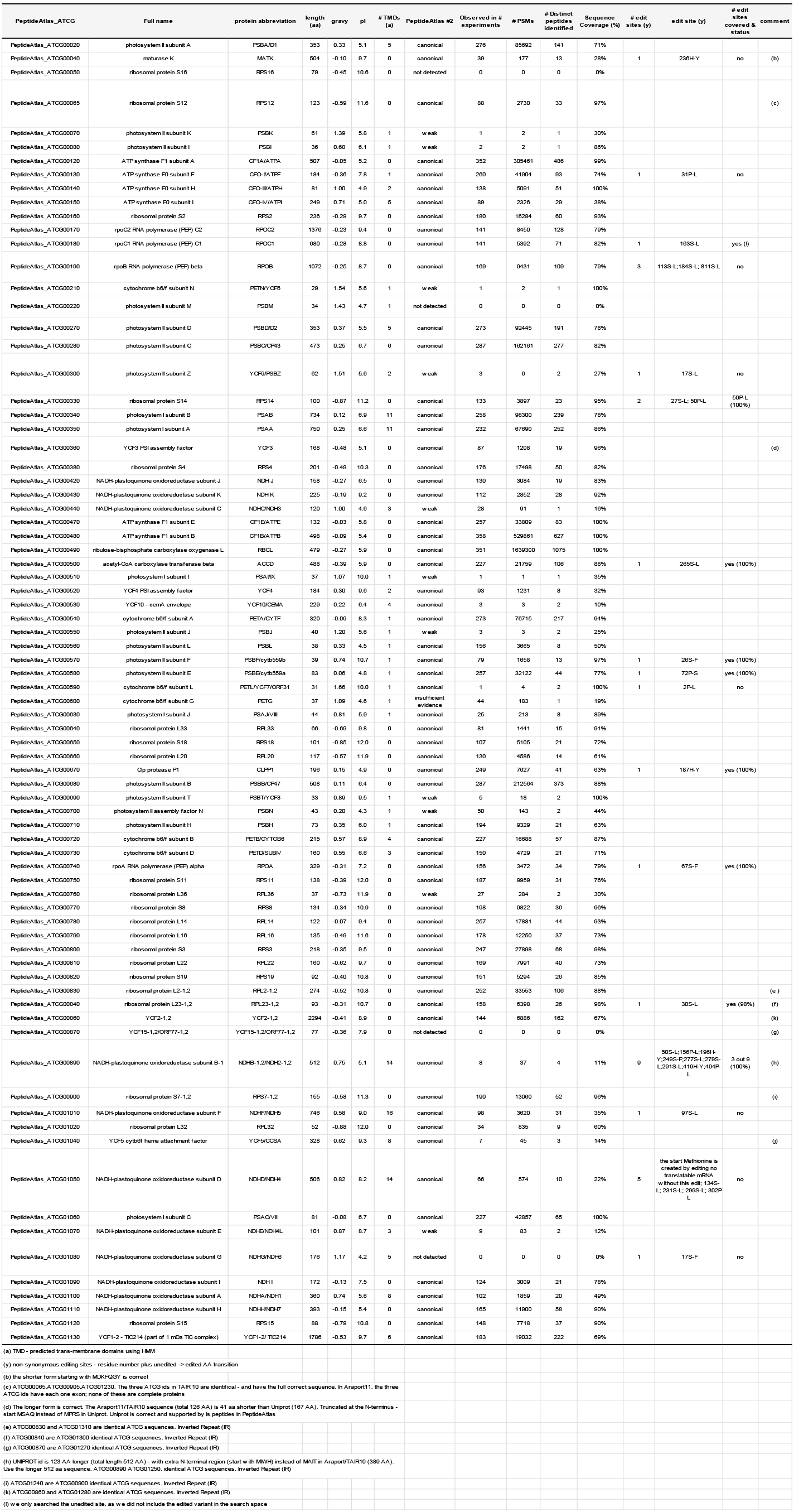
Consensus annotation, non-synonymous editing sites, protein identification and editing status of the plastid-encoded proteins based on Arabidopsis PeptideAtlas build 2.

Table 2 summarizes the mitochondrial-encoded proteins with their gene ATMG identifier, protein name and function, and editing events that impact the protein sequence. These identifiers and their unedited and (partially) edited amino acid sequence variants can be downloaded at Arabidopsis PeptideAtlas. This table is mostly based on mitochondrial genome sequences for Arabidopsis ecotype C24 and Col-0 (Unseld et al., 1997; Davila et al., 2011), the extensive body of mitochondrial literature, *e.g*. (Rao et al., 2016; Planchard et al., 2018; Moller et al., 2021)), and the recent study in which publicly available Illumina MiSeq data were used to perform *de novo* assembly of the Arabidopsis Col-0 mitochondrial genome, followed by additional verifications against other experimental DNA sequence data (Sloan et al., 2018). There are 32 unique mitochondrial-encoded protein sequences and three ORFs (Table 2). Complete protein-coding transcripts for mitochondrial-encoded NDH1,2,5 are each generated by trans-splicing of individual exons. Each of these exons have ATMG identifiers in Araport11/TAIR 10 but these three proteins should each be represented by one ATMG identifier; for simplicity we selected the identifiers with the lowest number (Table 2). These 32 mitochondrial-encoded proteins are nine subunits of the NAD complex (complex I), one subunit of complex III, three subunits of complex IV, six subunits of the ATP synthase complex (complex V), five proteins involved in cytochrome biogenesis, seven ribosomal proteins, one maturase. In addition, there are the tentative proteins TAT ORFX/TATC (candidate transporter protein), ORF114 and ORF240A. All mitochondrial protein coding RNAs, except COX1 and ORF114, are predicted to undergo RNA editing that lead to one or more amino acid change. There are in total predicted to be 420 edits that would result in a change in amino acid. Based on RNA data, 57 of these editing sites are considered low frequency sites (Table 2); essentially this means when sequencing cDNA libraries, these mRNAs are mostly found in their unedited form. These edit sites are based on information from (Bentolila et al., 2013; Sloan et al., 2018) and other publications, as well as from communications with the scientific community with expertise in plant mitochondrial editing (see Acknowledgements).

**Table 2.**
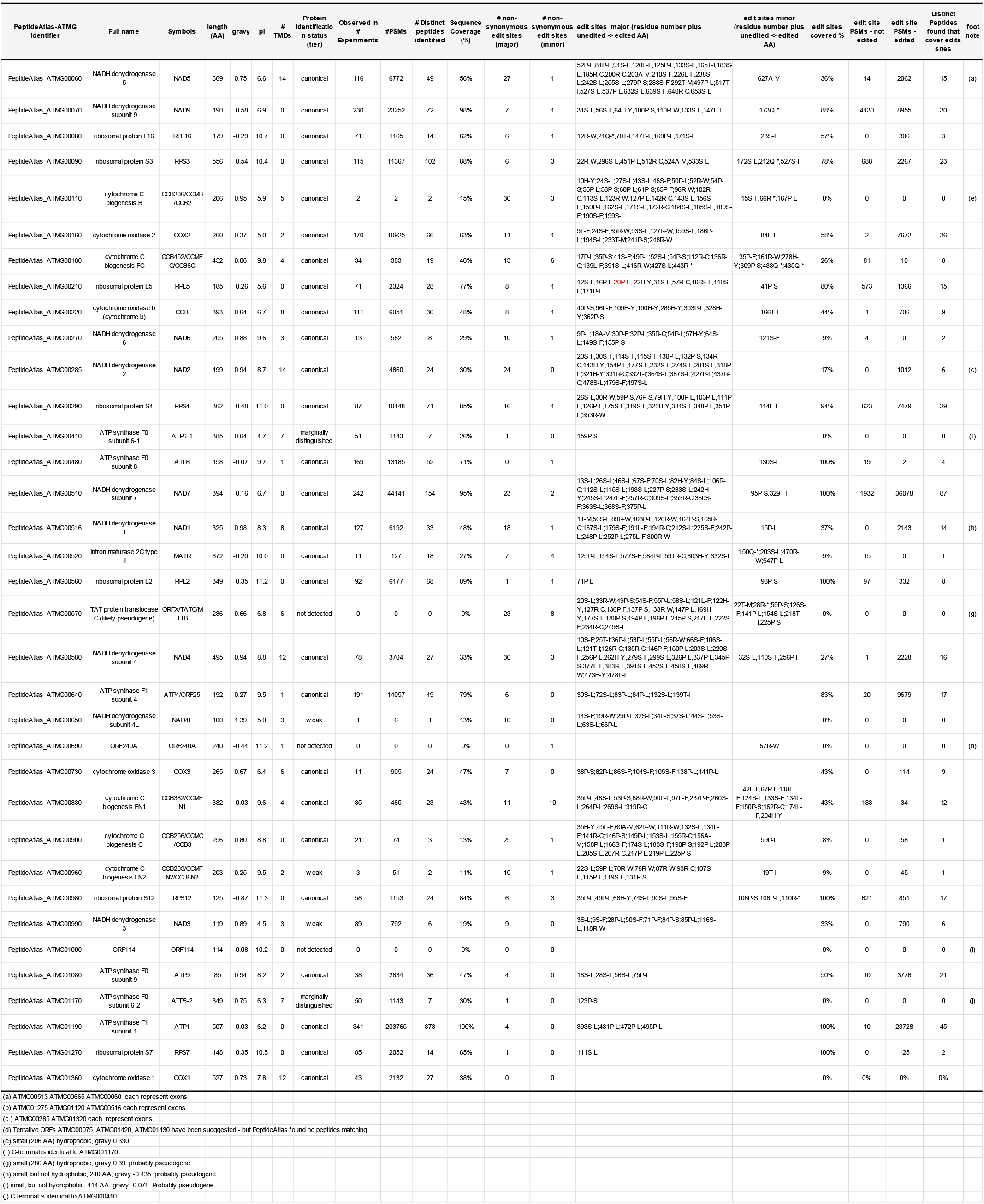
Consensus annotation, non-synonymous editing sites, protein identification and editing status of mitochondrial-encoded proteins based on Arabidopsis PeptideAtlas build 2.

### Detection of the predicted unedited and edited plastid proteome

We detected 63 plastid-encoded proteins at the most confident (canonical) level, as well as 12 proteins at lower confidence levels (tiers), whereas three proteins (NDHG/NDH6, PSBM and RPS16) and ORF YCF15 were not detected, *i.e.* there were no matched MSMS spectra (PSMs) that passed our confidence threshold. The lower confidence level identifications fall in the two tiers, *i.e.* insufficient evidence (one or one uniquely mapping peptides but none reach 9 residues in length) or weak (at least one uniquely mapping peptide of ≥ 9 residues in length, but otherwise does not meet our criteria for canonical). The system of different confidence tiers was explained in detail in (van Wijk et al., 2021). Protein sequence coverage (by mapped peptides) of these identified proteins ranged from 10% (YCF10/CEMA) to 100% (Table 1). As expected, RBCL was identified with the highest number of PSMs (1.6 million), followed by CF1β and CF1α with 0.5 and 0.3 million PSMs, respectively (Table 1). The three undetected proteins and the proteins assigned ‘weak’ identifications are either very hydrophobic (gravy indices > +1, and/or small (<100 aa). RPS16 is a 79 aa basic protein (isoelectric point 10.57) due the presence of a high number of K and R residues; such high basicity is quite common for ribosomal proteins. We were surprised by its lack of identification because tryptic digestion of the protein should result in several tryptic peptides of at least 7 aa residues. It very likely that RPS16 is in fact a pseudogene in Arabidopsis. Widespread pseudogenization of *RPS16* in the angiosperm chloroplast genomes via the loss of its splicing capacity was described, even when the *RPS16* encoded in the chloroplast genome is transcriptionally active (Ueda et al., 2008; Roy et al., 2010). It was suggested that the chloroplast-encoded RPS16 in mono– and dicotyledonous plants has been substituted by the product of nuclear-encoded RPS16 which was transferred from the mitochondria to the nucleus before the early divergence of angiosperms (Ueda et al., 2008; Roy et al., 2010). Arabidopsis Col-0 has two nuclear-encoded RPS16 homologs (AT4G34621 and AT5G56940; RPS16-1 and –2, respectively) of which AT4G34621 was shown to be an essential gene (Tsugeki et al., 1996). Based on confocal microscopy with fluorescent reporter constructs, it was suggested that the RPS16 homologs have dual-localization signals for plastids and mitochondria (Ueda et al., 2008). Arabidopsis PeptideAtlas and PPDB shows that RPS16-1 accumulates at much higher levels than RPS16-2 and inspection of the metadata suggest that RPS16-1 localizes to chloroplasts whereas RPS16-2 localizes to mitochondria. The 176 aa NDHG/NDH6 (gravy 1.17) was not identified because it has five predicted transmembrane domains and only on lysine (K128) and one arginine (R175); Hence trypsin digestion does not result in any peptides suitable for detection by MS. It was shown that its translation is under control of PGR3 and it is not a pseudogene (Rojas et al., 2018; Higashi et al., 2021). PSBM is a 34 aa protein with one TMD and contains only one lysine (K28) and no arginine. Trypsin digestion will result in a 28 aa N-terminal peptide spanning the TMD and a six aa C-terminal peptide which is below the minimal length for protein identification; this explains the lack of observation. PSBN is located in the stroma lamellae and is involved in biogenesis of PSII, but is not a PSII subunit itself (Plochinger et al., 2016).

Chloroplast RNA editing (or lack of editing) at the protein level may be detected in the form of (un)edited peptides; these peptides must have at least 7 amino acids and a decent propensity to be ionized in the source of the mass spectrometer. In total, after manual validation, we observed peptides covering 10 of the 32 editing sites across 8 plastid-encoded proteins (Table 3). The level of experimental support for these editing sites is determined by the number of distinct peptides and the number of observations (PSMs). For the highly observed sites, 72P-S in ATCG00580.1 (cytb559α) and 30S-L in ATCG00840.1 (RPL23A) these unedited sites were only at 0.2% and 2% frequency, respectively. The RPOC1 editing site (163S-L) was detected with only 8 PSMs and only one distinct peptide; but none showed the edited form. This is consistent with the relatively low levels of RNA editing (∼10%) observed at the transcript level in wt Arabidopsis plants (Chateigner-Boutin et al., 2010). For the other six editing sites, only the edited form was detected. Figure 1 displays some representative spectra for both the unedited and edited forms for these two cases (cytb559α and RPL23A). These four PSMs are of extremely high quality and provide unequivocal evidence of both edited and unedited forms. To evaluate possible biological significance for these partial edits, we looked at the metadata for particular features of the samples *(e.g.* non-photosynthetic tissue). However, no obvious trends were found, and we hypothesize that some low level of missed editing is inevitable in plastids and it is not detrimental to their overall successful function.

**Figure 1.**
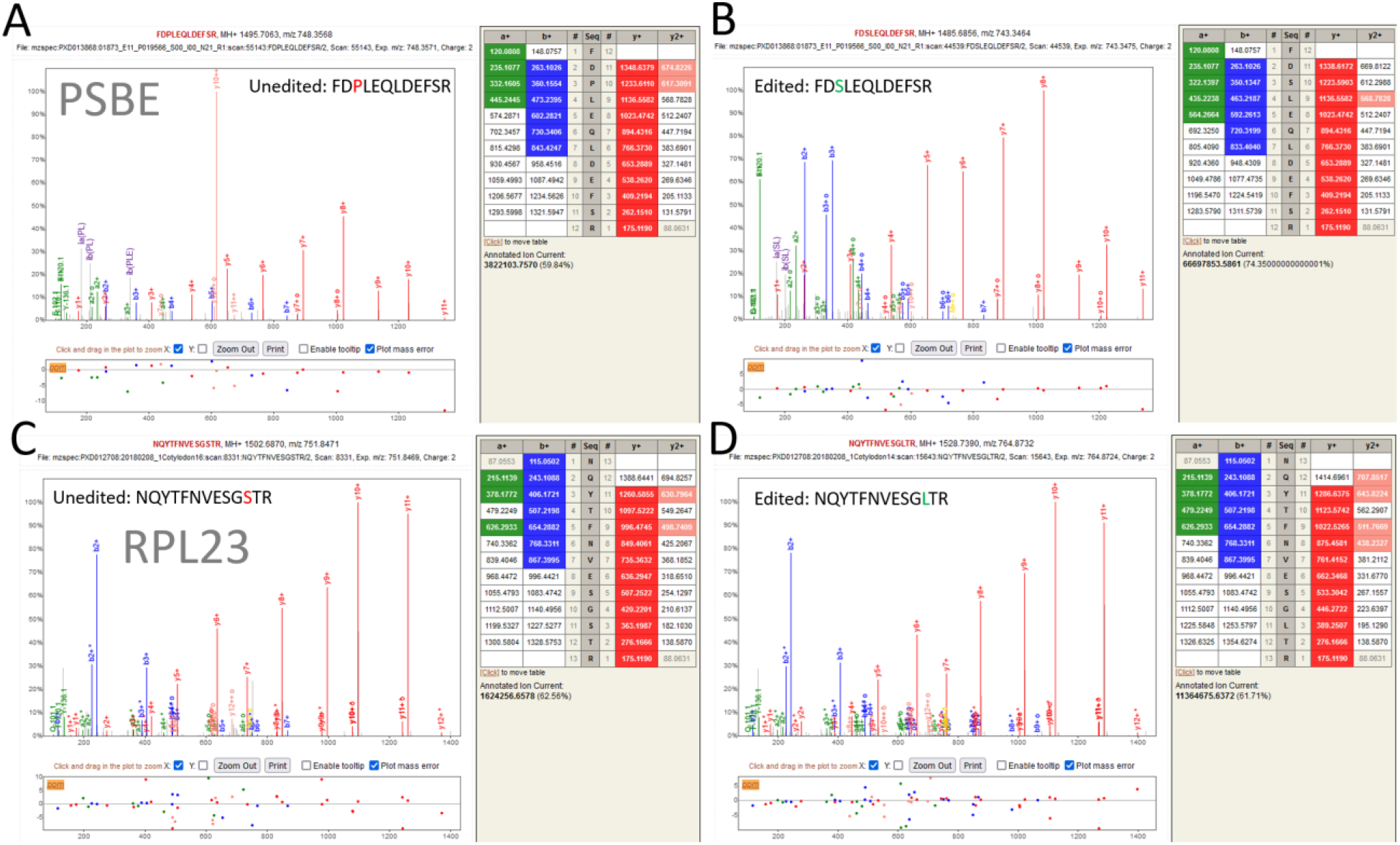
Representative spectra for unedited and edited forms of cytb599α (PSBE) and RPL23-1,2. A) the unedited form FDPLEQLDEFSR of 72P/S of cytb599α. Note the very strong y10++ ion indicative of a proline. B) the edited form of FDSLEQLDEFSR of 72P/S of cytb599α with a visible but muted y10++ ion. C) the unedited form NQYTFNVESGSTR of 30S/L of RPL23. Detection of nearly all y ions provides ample evidence of the unedited form. D) the edited form NQYTFNVESGSTR of 30S/L of RPL23 with an equivalently perfect ladder of y-ions.

**Table 3.**
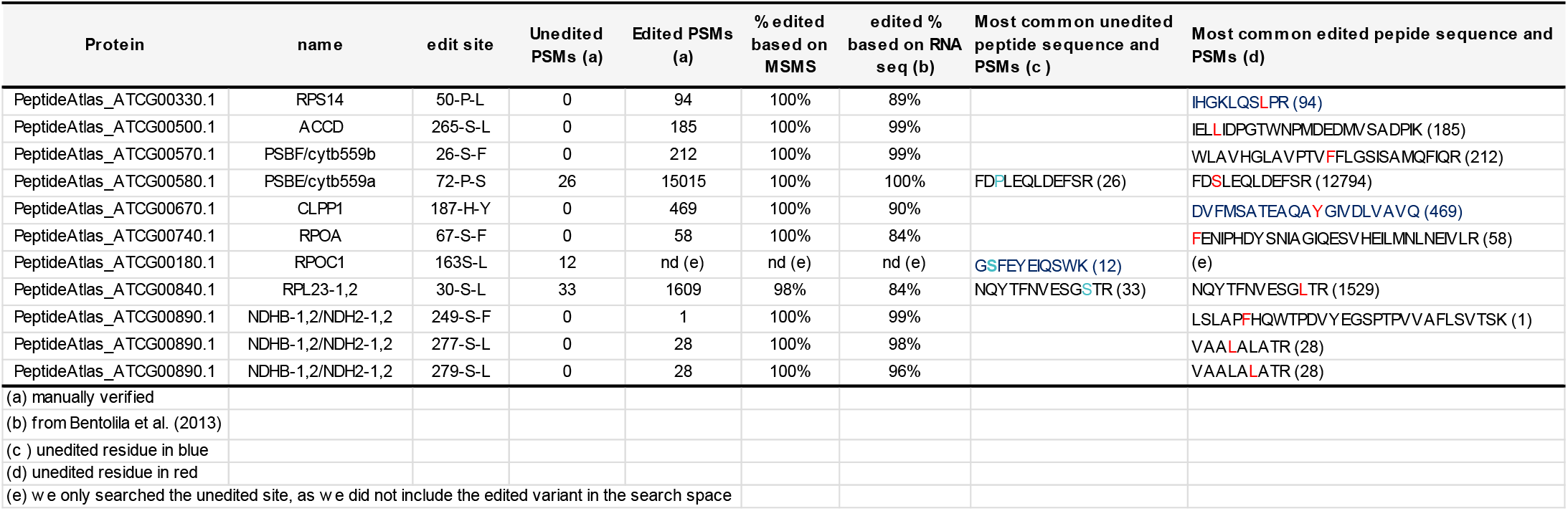
Verified protein editing status of 10 editing sites in eight plastid-encoded proteins based manual evaluation of the 2^nd^ release of the Arabidopsis PeptideAtlas.

### Detection of the predicted unedited and edited mitochondrial proteome

We detected all 32 mitochondrial-encoded proteins, but none of the three ORFs, with 27 proteins at the most confident (canonical) level and five at lower confidence levels (Table 2). Sequence coverage ranged from 11% (CCB203/CCMFN2/CCB6N2) to 100% (ATP1), and matched PSMs ranged from two (CCB206/CCMB/CCB2) to ∼0.2 million PSMs (ATP1). The five proteins identified at lower confidence levels are the pair ATP6-1,2 homologs (identical in their C-terminal 252 aa) with a few peptides that were able to distinguish between ATP6-1 and ATP6-2 (Table 2), NDH4L, NAD3 and CCB203/CCMFN2/CCB6N2. There were no MSMS spectra (PSMs) that mapped to the three ORFs (ORF114, ORF240A and ORFX/TATC). If translated, these three ORFs are of medium size (114, 240 and 206 aa) and not particularly hydrophobic (0.078, 0.435 and 0.39) with ample tryptic cleavage sites, and thus their physicochemical properties do not interfere with detection by MSMS. Therefore, the complete lack of observation (matched spectra) suggests that these three ORFS are in fact pseudogenes.

The mitochondrial intron maturase 2C type II (matR) (ATMG00520.1 and P93307) is annotated in several sources as having a non-AUG start site. As shown in figure 2A, sequences from UniProtKB (P9907), TAIR10 (TAIR10_ATMG00520.1), Araport11 (Araport11_ATMG00520.1), and the PeptideAtlas consensus (PeptideAtlas_ATMG00520.1) all have a non-methionine start site. Sloan et al. (Sloan et al., 2018) annotated a later start site at the first methionine 17 residues further downstream, represented in UniProtKB with A0A2P2CLH3 and RefSeq as YP_009472117.2. UniProtKB cites (Sloan et al., 2018) for this entry. Obtaining MS evidence of the methionine start site is difficult on account of the closely space lysine and arginine residues that follow the methionine, and that part of the sequence is indeed not represented in PeptideAtlas. There is one peptide detected in PeptideAtlas (FRPLTVVLPIEK) that appears to support the annotated non-AUG start site and N-terminus (GLKFRPFRPLTVVLPIEK). This tryptic peptide (FRPLTVVLPIEK) is supported by six PSMs that pass our threshold in four separate samples from two datasets (PXD010324 and PXD013868). Most of these spectra have a signal-to-noise ratio less than 100 and are not very impressive, but one higher quality spectra shown in Figure 2B appears to provide quite compelling evidence for the detection of FRPLTVVLPIEK. This spectrum (USI mzspec:PXD013868:01874_B05_P019568_S00_I00_N02_R1:scan:49487:FRPLTVVLPIEK/3) has nearly all major peaks explained, albeit with substantial internal fragmentation. This peptide does not map to any other entries in our comprehensive database, even considering I/L substitution. There is growing evidence that many proteins sometimes or often begin translation at a non-AUG start site (Cao and Slavoff, 2020). However, glycine is an unusual start residue, whereas GUG (valine) and UUG (leucine) are the most common after AUG (methionine) (Belinky et al., 2017). It may be that the more likely start site is the leucine following the glycine (hence: N-terminus is LKFRPFRPLTVVLPIEK).

**Figure 2.**
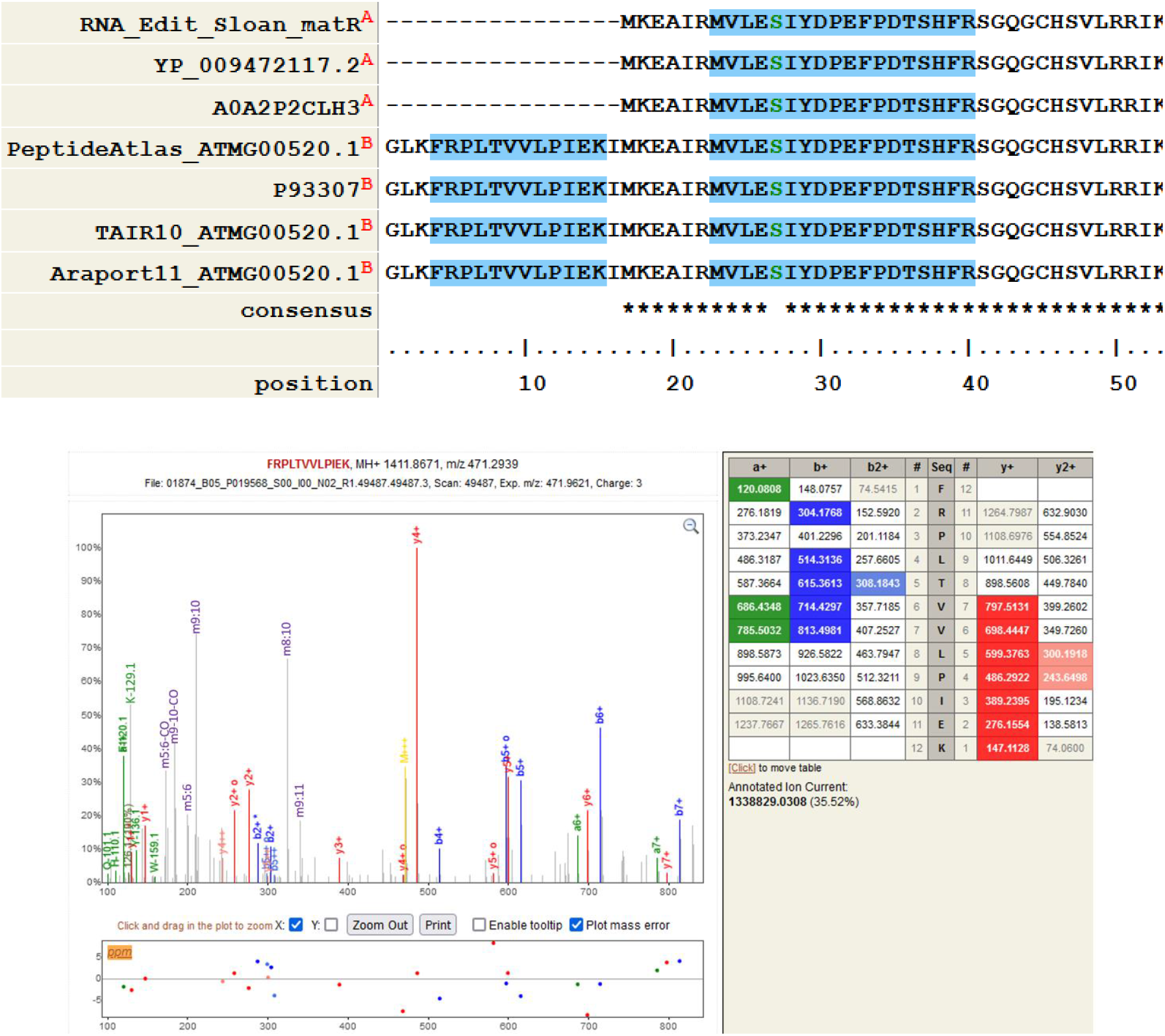
A sequence alignment of mitochondrial matR in several different sources. UniProtKB’s primary entry, TAIR10, and Araport11 all maintain an early non-AUG start site, while UniProtKB alternative entry A0A2P2CLH3 and RefSeq have a later start site at the first methionine, citing Sloan et al. Sequence with a blue shading is detected in PeptideAtlas, while white background is not detected. The serine is green at position 27 is a subject of a potential variant, not detected in PeptideAtlas. B) mzspec:PXD013868:01874_B05_P019568_S00_I00_N02_R1:scan:49487:FRPLTVVLPIEK/3 using the Lorikeet spectrum viewer that shows compelling evidence for the detection of FRPLTVVLPIEK, a peptide that only maps to a potential early start site of matR. The spectrum may be examined at http://proteomecentral.proteomexchange.org/usi/ using the above USI. The internal fragmentation ions (*e.g.* m9:10 indicates a b-type ion of residues 9-10, i.e. PI) are not natively labeled in the spectrum viewer (Lorikeet) and added manually above.

There are in total 420 predicted non-synonymous edits across the mitochondrial-encoded proteins. Based on RNA data, 57 editing sites are considered low frequency sites (Table 2). In total we observed 157 of the editing sites (Tables 2,4). We applied the same criteria and manual evaluation of low frequency events as for the plastid-encoded proteins. Manual evaluation uncovered several incorrect assignments for the S/L editing site with assignment of formylated S instead of L, the edited form. This was due to a software assignment error in selection of the correct isotope, which should be the monoisotopic form of the selected precursor ion (*i.e*. all carbon atoms are C12) but instead the precursor ion with one C13 was selected. Figure 3 shows an example of a peptide for NAD7 that appears to provide evidence of unedited protein, but the explanation provided by edited form is better, and the most likely explanation is that the instrument-selected precursor mass belongs to a co-fragmented contamination precursor, rather than the proposed explanation. This demonstrates one of challenges in making unedited/edited assignments, and why manual inspection of some assignments is required.

**Figure 3.**
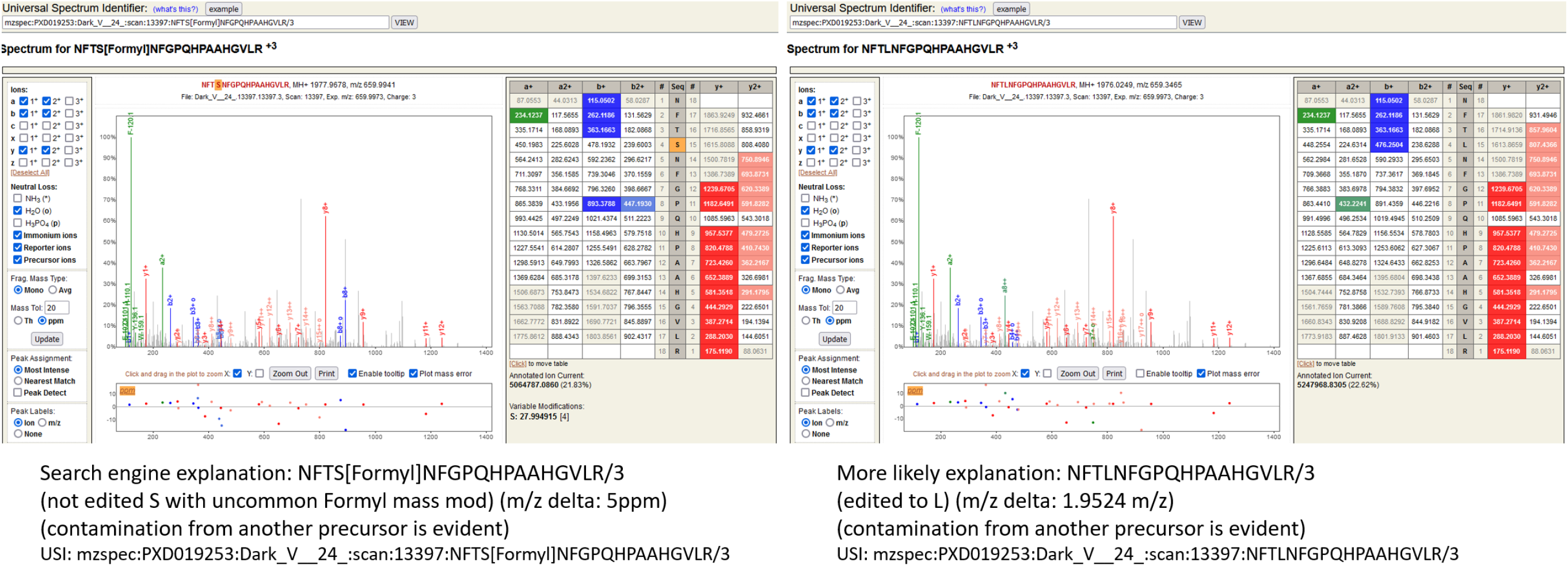
Example of likely unedited/edited conflation. The left panel shows the search engine result, which matches the spectrum to an unedited peptide, but with the uncommon formylation mass modification (+27.9949) on the unedited serine. The precursor m/z delta is only 5ppm, which is within our search tolerance. Although some unlabeled gray peaks can be explained by internal fragmentation and imperfect annotation, major unexplained peaks at 732 and 892 are clearly contamination by another co-fragmented precursor. The b8 annotation is spurious as it is on the wrong isotope and completely unexpected on the wrong side of a proline. The right panel shows a more likely explanation, after manual inspection. The S[Formyl] is replaced by the expected editing result, an L. This allows both the b+ and y++ series to extend further in both directions. The spurious b8 in the left explanation disappears. However, the precursor m/z is now way off at 1.9524 m/z, well outside our search tolerance. Attempts to rectify this with deamidation do not improve the fragmentation explanations. The most likely explanation is that the instrument-selected precursor mass belongs to the contamination precursor, rather than the proposed explanation. This demonstrates an example of potential hazards in making unedited/edited assignments, and why manual inspection of some assignments is required. The spectrum and potential explanations can be explored by the reader using the provided USIs via the ProteomeXchange USI page at http://proteomecentral.proteomexchange.org/usi/

The final validated editing site observation counts for mitochondrial-encoded proteins are provided in Table 4 organized by specific editing site. Figure 4 displays these results for the 25 proteins with at least one editing site observed with at least 5 PSMs. Nearly all sites have nearly complete edits at 95%-100% or non-edits 0%-10%, with the single exception of RPL5, with an edited/unedited ratio of 0.6. In case of one minor frequency site based on RNA (ATMG00830 124S/L with 5/0 edited/unedited), we observed high frequency at the peptide level. Importantly, for all other 20 edit sites that we observed with low editing frequency were indeed assigned as low frequency site at the RNA level (Sloan et al., 2018). Conversely, all sites that we observed to be edited at high frequency were also assigned as high frequency editing sites at the RNA level.

**Figure 4.**
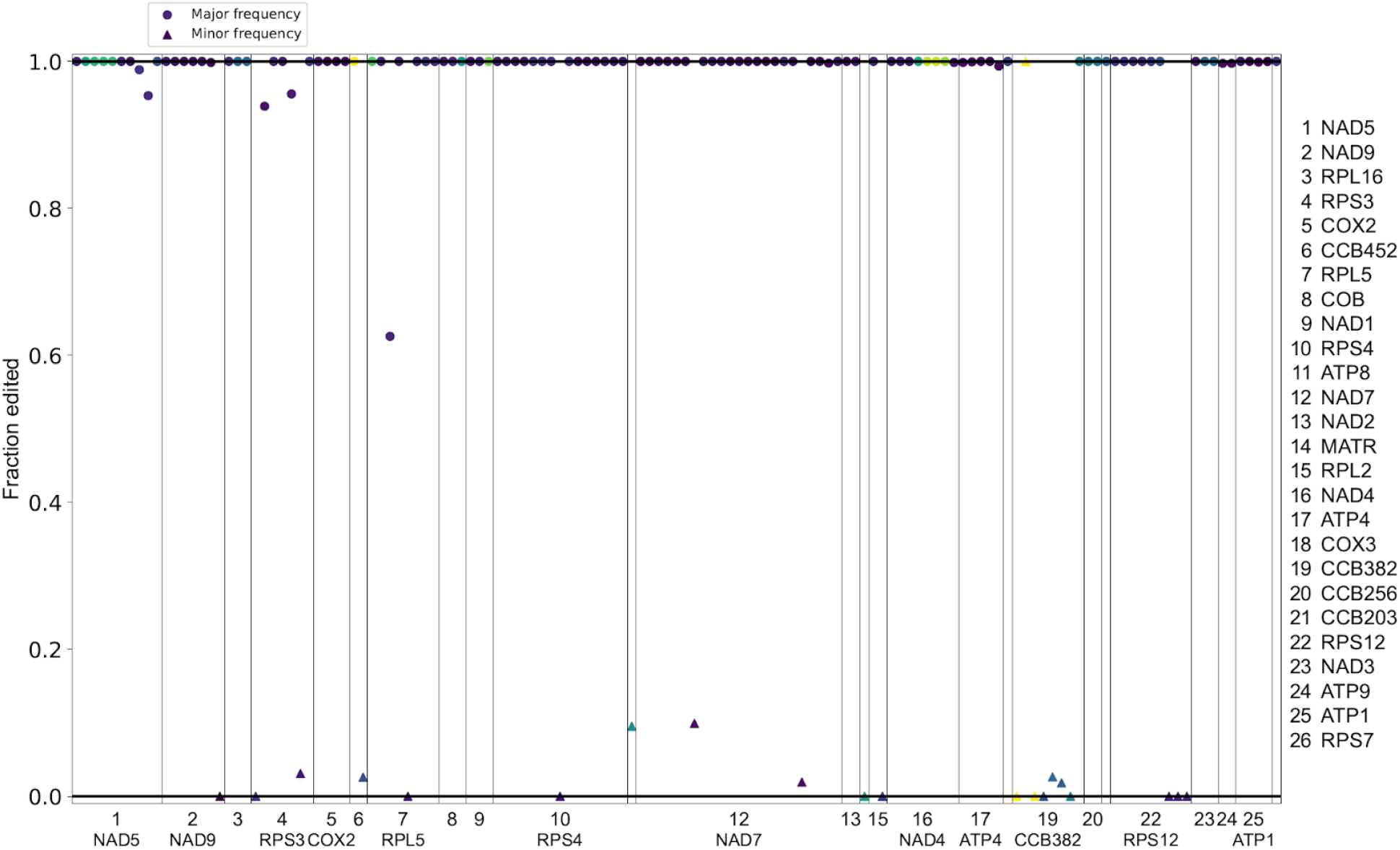
Overview of mitochondrial editing sites based on Table 4 for the 26 proteins with at least one editing site with at least 5 observations. High frequency edits (based on RNA) are depicted with circles and low-frequency edits (based on RNA) with triangles. Vertical lines separate the 25 proteins, which are numbered at the bottom and labeled on the right. Observed sites at higher frequencies are darker in color, while sites with low numbers of observations are shown with a lighter color.

**Table 4.**
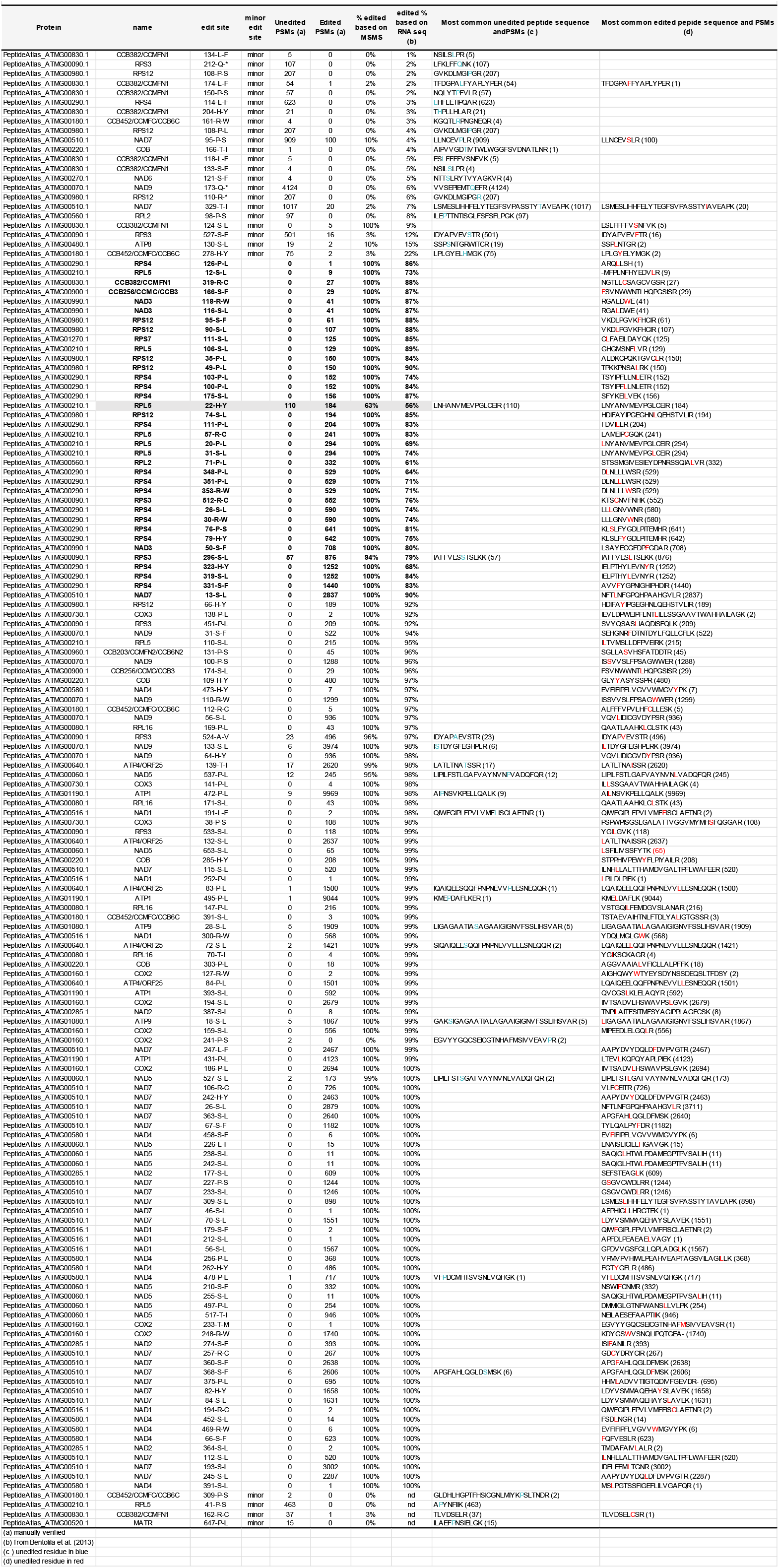
Verified protein editing status of 156 editing sites in 27 mitochondrial-encoded proteins based manual evaluation of the 2^nd^ release of the Arabidopsis PeptideAtlas.

### Frequencies and observations of the different type on edits at the amino acid level

RNA editing across plastids and mitochondria leads to 14 possible amino acid changes (accounting for these 452 total possible edits) including two types of RNA edits (Q->stop; R-> stop) that lead to introduction of a stop codon (six and four sites, respectively) (Table 5). The predicted S to L edit is the most frequent amino acid change (117 times), whereas the T to M edit is the least frequent (in just five cases). Across plastid and mitochondrial proteins, we observed all types of edits at the peptide level with the exception of the P to F change, and the stop codon introductions (Tables 4 and 5). Evaluation of an introduction of a stop codon was challenging as this requires the identification of a diagnostic C-terminal peptide (and these stops would lead to truncation, rather than extension). We did not observed peptides that support new stop codons that result in such truncations. In 166 out of these 452 cases we did obtain peptide coverage of the editing site, such that we could potentially determine if an edit did indeed occur (Table 5). Because sequence identification by MSMS is based on predicted mass for each residue, amino acid changes are detected as a change in mass. Therefore, we calculate the predicted delta mass for each of the 12 changes (Table 5) and they range from 10.02073 to 60.03638 Da, which is well within the mass accuracy for modern mass spectrometers. Only in case of the S-L and P-L edit did the delta mass have some similarity to a common post-translational modification. The S-L transition corresponds to 26.05203 Da which is somewhat close (delta is 1.94288 Da) to formylation (27.99491 Da) and the P-L transition corresponds to 16.0313 Da which is somewhat close (delta is 0.03639 Da) to oxidation (15.99491 Da). However, these are easily distinguishable with the mass accuracy of current mass spectrometers and should not result in false positives. However, as we discussed above and figure 3, a few cases of misassignment of S to formylated S, rather than S to L were observed due to simultaneous fragmentation of two peptides and incorrect selection of the mono-isotopic peptide peak.

**Table 5.** Summary of consensus editing types at the amino acid level (*i.e*. consequences of edits) and their frequencies across plastid– and mitochondrial encoded proteins.

| Unedited->Edited | # searched edit sites (major & minor) | # Detected major sites (edited or unedited) | # Detected minor edit sites (edited or unedited) | % Detected | # PSMs unedited major | # PSMs edited major | # PSMs unedited minor | # PSMs edited minor | Mass of unedited AA (Da) | Mass of edited AA (Da) | Delta Mass (Da) |
| --- | --- | --- | --- | --- | --- | --- | --- | --- | --- | --- | --- |
| A-V | 6 | 1 | 0 | 17% | 23 | 496 | 0 | 0 | 71.03711 | 99.06841 | -28.0313 |
| H-Y | 27 | 12 | 2 | 52% | 110 | 8974 | 96 | 2 | 137.05891 | 163.06333 | -26.00442 |
| L-F | 21 | 3 | 4 | 33% | 0 | 2484 | 687 | 1 | 113.08406 | 147.06841 | -33.98435 |
| P-F | 8 | 0 | 0 | 0% | 0 | 0 | 0 | 0 | 97.05276 | 147.06841 | -50.01565 |
| P-L | 92 | 29 | 2 | 34% | 24 | 34190 | 222 | 0 | 97.05276 | 87.03203 | 10.02073 |
| P-S | 40 | 7 | 6 | 33% | 28 | 18341 | 1735 | 100 | 97.05276 | 113.08406 | -16.0313 |
| Q * | 6 | 0 | 2 | 33% | 0 | 0 | 4231 | 0 | 128.05858 | n/a | 128.05858 |
| R * | 4 | 0 | 1 | 25% | 0 | 0 | 207 | 0 | 156.10111 | n/a | 156.10111 |
| R-C | 30 | 7 | 1 | 27% | 0 | 1820 | 37 | 1 | 156.10111 | 103.00919 | 53.09192 |
| R-W | 30 | 8 | 1 | 30% | 0 | 4775 | 4 | 0 | 156.10111 | 186.07931 | -29.9782 |
| S-F | 55 | 16 | 3 | 35% | 6 | 10813 | 509 | 16 | 87.03203 | 147.06841 | -60.03638 |
| S-L | 117 | 54 | 2 | 48% | 122 | 45087 | 19 | 7 | 87.03203 | 113.08406 | -26.05203 |
| T-I | 11 | 3 | 2 | 45% | 17 | 3570 | 1018 | 20 | 101.04768 | 113.08406 | -12.03638 |
| T-M | 5 | 1 | 0 | 20% | 0 | 1 | 0 | 0 | 101.04768 | 131.04049 | -29.99281 |
| **Totals** | **452** | **141** | **26** |  | **330** | **130551** | **8765** | **147** |  |  |  |

One other consideration that might lead to biases in detection is the change in physicochemical properties of the amino acid of the unedited and edited position, possibly biasing the detectability by MSMS. In particular, the transition from R to W, R to C and H to Y, results in the loss of positive charge and increase in hydrophobicity, as well the loss of a tryptic cleavage site (since trypsin cleaves after R and K). The loss of the tryptic cleavage site can however be beneficial for improved coverage of the predicted edit site as a larger peptide is generated. We mapped a significant portion (25-55%) for all types of RNA editing sites, with the exception of the two P-F transitions (Table 5). Therefore, the editing frequencies of plastid and mitochondrial proteins were therefore unbiased for editing frequencies for the different types. Finally, the MSMS results mapped 26 out of 56 RNA-seq-based minor frequency sites (46%) compared to 140 out 395 RNA-seq-based high frequency sites (35%), indicating that the peptide analysis was unbiased for its assessment of low (minor) vs high frequency edits.

### Post-translational modifications of the plastid– and mitochondrial-encoded proteomes

Like nuclear-encoded proteins, the organelle-encoded proteins can undergo several types of *in planta* post-translational modifications (PTMs). We specifically searched for N-terminal acetylation for all proteins, and for lysine acetylation, STY phosphorylation and lysine ubiquitination in case of experimental data sets that were deliberately affinity enriched for these PTMs (van Wijk et al., 2023). A sophisticated PTM viewer in PeptideAtlas allows detailed examination of these PTMs, including direct links to all spectral matches and we recommend using the PeptideAtlas to evaluate specific PTM sites if these are of particular interest to the reader. PTM identification rates strongly depend on the confidence level (minimal probability threshold) of PTM assignment. Here we used localization probability P≥0.95 from PTMProphet (Shteynberg et al., 2019) for each PTM, and also required at least 3 PSMs for a specific PTM at a specific residue to be included in the summary (Supplemental Data Set S1 for the data on all four PTMs). In general, higher numbers of repeat observations (PSMs) for a specific PTM at a residue improve the reliability of the assignment. Conversely, peptides with high PSM counts (*e.g.* hundreds or more) for which the vast majority (*e.g.* >90%) of peptide do not have a reported PTM at P>0.95, are possibly false discoveries. We evaluated the results for false positives and possible pitfalls in various ways, including spot checking matched spectra and proteins to which PTMs were mapped. Supplemental Data S1 provides these details and also provide direct links to peptide and spectral URLs (within the PeptideAtlas DB).

The chloroplast acetylation machinery in Arabidopsis consists of eight General control non-repressible 5 (GCN5)-related N-acetyltransferase (GNAT)-family enzymes that catalyze both N-terminal and lysine acetylation of proteins, whereas no mitochondrial localized protein acetyltransferases are known (Pozoga et al., 2022). N-terminal acetylation (NTA) at position 1 (the start Methionine) or 2 was observed for 17 plastid proteins, but not for any of the mitochondrial-encoded proteins (Supplemental Data Set S1). This is consistent with previous reports that chloroplast-encoded proteins undergo frequent NTA (here observed at residues A, S, T, V, I, R) (Zybailov et al., 2008; Dinh et al., 2015; Rowland et al., 2015; Bienvenut et al., 2020; Willems et al., 2021) but mitochondrial proteins do not (Huang et al., 2009). For a few of these chloroplast-encoded proteins we observed NTA at both position 1 and 2 (PSAA, CF1A, CF1B, D1) but NTA at position 2 always had far more PSMs. For CF1A, CF1B, RBCL, and CP47 the search also identified NTAs much farther down-stream of the N-termini. However, in nearly all cases these likely represent false positives since most of these peptides were observed without NTA (*i.e.* the frequency of assigned NTA for these observed peptides was very low). We observed lysine side-chain acetylation of only two mitochondrial proteins (ATP1 and NAD1) (4 sites), but 15 chloroplast proteins belonging to the ATP-synthase (CF1α,β, CFO-I,III), the cytb6f complex (cytf and cytb6), Photosystem I (PsaA,B), Photosystem II (CP43,CP47,D2,PsbH), TIC214 and RBCL (66 sites). The number of PSMs per K-acetyl site that passed the p-value thresholds range from 3 to 159; higher number of PSMs should associate with higher overall confidence of significance of the observation. However, some PTM sites were observed with very high number of PSMs that pass the PSM FDR quality threshold with a PTM at that site, irrespective of the PTM localization probability (Supplemental Data Set S1). The ratio between PSMs that pass the thresholds and total PSMs can be reflection of FDR, and PTM sites with low ratios should be treated with reservation.

We observed phosphorylation of 12 plastid-encoded proteins and just one mitochondrial protein (RPS3) (S-112) (Supplemental Data Set S1). The phosphorylated plastid proteins include RBCL, ribosomal RSP7/14, YCF1, PSAA, three Photosystem II core proteins (CP47, PSBH, PSBL), four ATP-synthase subunits. The number of PSMs for most of these p-sites was relatively low, especially compared to the high # PSMs for these peptides from non-phosphorylation data sets (which were not searched for p-peptides). p-sites with only few PSMs are therefore very rare events especially considering the large number of spectra searched; however, several p-sites (RBCL-330T and RPS7-S93) were also observed in prior, carefully evaluated data sets (Schonberg et al., 2017).

Based on the two large scale ubiquitination studies included in build 2 (Walton et al., 2016; Grubb et al., 2021), we observed only ubiquitination of chloroplast RPS7 (note that we observed ubiquitination for more than 1000 nuclear-encoded proteins (van Wijk et al., 2023). This UBI site in RPS7 (K13) was only observed in the data from (Walton et al., 2016) and with only 3 PSMs; the significance of this observation is not clear but could represent an unintended alkylation which results in an identical mass. The lack of ubiquitination for organelle-encoded proteins is consistent with our re-analysis of a very recent ubiquitination study that was otherwise not included in the 2^nd^ Arabidopsis PeptideAtlas release (van Wijk et al., 2023).

## DISCUSSION

### Consensus sequences, identification and coverage of the plastid mitochondrial-encoded proteomes

Originally sources reported around 88 protein-coding plastid genes and 31-33 protein-coding mitochondrial genes in Arabidopsis (Unseld et al., 1997; Sato et al., 1999; Sloan et al., 2018). Unfortunately, several of the last iterations of the Arabidopsis community genome annotations (including TAIR10 and Araport11), include 122 mitochondrial-encoded protein identifiers (ATMG). It is not entirely clear why these additional identifiers and sequences were introduced. Furthermore, both plastid and mitochondrial cDNAs are generated by transplicing; at the protein level, these individual exons are not stable protein products and the proteins from by multiple spliced transcripts should be represented by a single protein identifier. Finally, the plastid genome includes 2 inverted repeats mostly encoding for identical protein products (the exception is the non-functional and truncated ATCG01000). To better determine the accumulation of the plastid– and mitochondrial proteins it is therefore imperative that a non-redundant set of identifiers of the full-length protein sequences is available. With help of several experts in the international research community, this study assembled this non-redundant set of proteins, and also resolved several conflicting start and stop sites (Tables 1 and 2). Based on the systematic reprocessing and database search of >250 million raw MSMS spectra as part of the new release of Arabidopsis PeptideAtlas, we confirmed accumulation of all but three of the plastid-encoded proteins (RPS16, PSBM, NDHG/NDH6), and all but three of the mitochondrial-encoded proteins (NAD4L; CCB203/CCMFN2/CCB6N2; NAD3). We conclude that plastid RPS16 is likely a pseudogene, and that plastid ORF77/YCF15 as well as mitochondrial ORF114 and ORF240A are also pseudogenes. Therefore, the plastid genome encodes for stable 77 proteins and the mitochondrial genome encodes for 32 stable proteins. However, due to RNA editing, most of the mitochondrial stable proteins and 17 plastid-encoded proteins are modified as compared to the unedited sequence.

To provide the research community this updated set of protein identifiers and their sequence variants, we provide several sequence datasets that can be used to *e.g.* simply obtain the most likely protein sequence variant for stable proteins, and/or for small– or large-scale proteomics studies, as follows:

**1.** protein ids and unedited sequences (with the exception of essential edits for start and stop codons that need to be applied to generate a protein) for the 79 plastid-encoded and 35 mitochondrial-encoded proteins and pseudogenes.
**2.** protein ids and sequences after application of editing for the 79 plastid-encoded and 35 mitochondrial-encoded proteins and pseudogenes; minor frequency edits are not applied but all high frequency edits are included.
**3.** proteins ids and the >10.000 sequence variants (see Materials and Methods) to allow for complete and exhaustive MSMS data base search of all possible edits and allow for partial editing (similar as we did in this study).
**4.** protein ids and their amino acid sequences with all possible variants supplied in PEFF format (https://www.psidev.info/peff) This PEFF format allows for encoding variants in the file. This is a PSI format that some MS search engines (*e.g.* Comet) can handle. It is essentially similar to FASTA, but one can encode “at position N in the reference sequence, the amino acid can also be X”. (this could also allow to include previously identified PTMs)

Finally, we anticipate that a consensus set of sequences and their editing variants will replace the current set of plastid– and mitochondrial proteins in the next and upcoming Arabidopsis genome annotation (TAIR12; tinyurl.com/Athalianav12) and can also be down-loaded from the Arabidopsis PeptideAtlas web site.

### Detection and abundance of plastid and mitochondrial-encoded proteins

The number of PSMs observed per plastid– or mitochondrial-encoded protein is an indication of detection efficiency and abundance. Protein abundance is the consequence of transcription rate, translational efficiency and protein turnover rate. Ribosome profiling, which determines the ribosome occupancy per gene showed that mitochondrial Arabidopsis transcripts greatly differ in ribosome association levels (Planchard et al., 2018). In particular, mRNAs encoding c-type cytochrome maturation (CCM) protein have particularly low ribosome densities when compared to mRNAs encoding respiratory chain subunits. The low translational activity of the five CCM genes is supported by the small number of PSMs observed (2-485) and compared to respiration chain subunits or ribosomal subunits (mostly >1000 and up to ∼204,000) (Table 2). Two of the mitochondrial ORFs undetected in PeptideAtlas, ORF114 and ORF240A, were also not revealed by ribosome profiling (also considering their favorable predicted protein properties for detection by MSMS), therefore can be dismissed as pseudogenes. The results concerning the mttB/TatC gene are more conflicting as no PSM were reported in PeptideAtlas, but a detection of translational activity by ribosome profiling may support its functionality (Planchard et al., 2018). Arabidopsis TatC lacks a classical start codon; however, an antibody raised against a TatC specific peptide was able to detect a faint band in isolated Arabidopsis mitochondria (Carrie et al., 2016). The binding was lost when the antibody was pre-incubated with the peptide, suggesting that the band was not a non-specific reaction. Furthermore, using immunoblotting and BN-SDS-PAGE TatC was found to colocalize with a newly identified TatB in the same high molecular weight complex. A definitive answer as to whether TatC is present in the Arabidopsis mitochondrial proteome would require the immunoprecipitation of the protein by the peptide antibody and analysis by MSMS of the immunoprecipitated proteins. Our study might also have revealed a non-canonical codon start for MATR, as 4 PSMs were detected for a 16 aa peptide upstream of the methionine start codon.

### Impact of RNA editing on the plastid and mitochondrial proteomes

This study is the first systematic MSMS-based analysis on the impact of RNA editing on the accumulated plastid and mitochondrial proteomes across many photosynthetic and non-photosynthetic tissues, developmental stages and (a)biotic conditions. Previously, only two different mitochondrial proteins (ATP6 and RPS12) encoded by partially edited transcripts were evaluated for the presence of peptides encoded by incompletely edited transcripts; it should be noted that this was in Petunia and maize, not Arabidopsis (Lu and Hanson, 1994; Lu et al., 1996; Phreaner et al., 1996). In the case of ATP6, protein from unedited transcripts was not detected despite the presence of partially edited transcripts on polysomes. (Lu and Hanson, 1994) took advantage of partial editing of a stop codon that shortens the protein by 13 aa, allowing production of an antibody against the unedited tail of ATP6. In contrast, antibodies raised to peptides specific to unedited portions of *rps12* transcripts could detect some accumulation of RPS12 encoded by incompletely edited transcripts (Lu and Hanson, 1994, 1996; Lu et al., 1996; Phreaner et al., 1996; Williams et al., 1998). The small number of proteins analyzed so far has prevented definitive conclusions about the possible role of partial editing in generating protein polymorphisms, especially in the plant mitochondrion. In mammalian systems such as for apolipoprotein B, C-to-U editing creates a stop codon so that two forms of the protein differing in size and translated from edited or unedited transcripts, are produced with different function and tissue-specificity (Davidson, 1993). Similarly, differential RNA editing in the human brain can alter the properties of glutamate receptors (Seeburg and Hartner, 2003). In contrast, the dogma in the plant organelle editing field is that RNA editing is a corrective mechanism allowing the production of functional proteins despite improper DNA sequencies (Small et al., 2020; Small et al., 2023). The importance of RNA editing for protein function is supported by the numerous editing mutants showing strong developmental defective phenotypes (Small et al., 2020). Furthermore, editing usually restores the occurrence of phylogenetically more conserved amino acids that are encoded by organisms that do not edit their RNA. However, the question of whether plant organelle editing results in the production of a pool of diverse proteins from differentially edited transcripts has never been experimentally addressed in an extensive way.

Figure 5 summarizes the quantitative relationship between frequency of recorded plastid and mitochondrial editing site detected at the level of RNA (by RNAseq – from (Bentolila et al., 2013) and proteins (only listing sites detected with at least 5 PSMs). This showed in case of minor editing sites (assignment based on RNAseq) these sites are also mostly only partially edited at the protein level; hence for these proteins, the lack of editing does not prevent proteins from accumulating. For major frequency editing sites, editing at the RNA level was between 60% and 100%; more than 30 of these sites editing was incomplete and well below 100% (Figure 5). However, editing at the protein level was nearly always 100% or close to 100%, except for RPS3 296-S-L at 94% (Figure 5).

**Figure 5.**
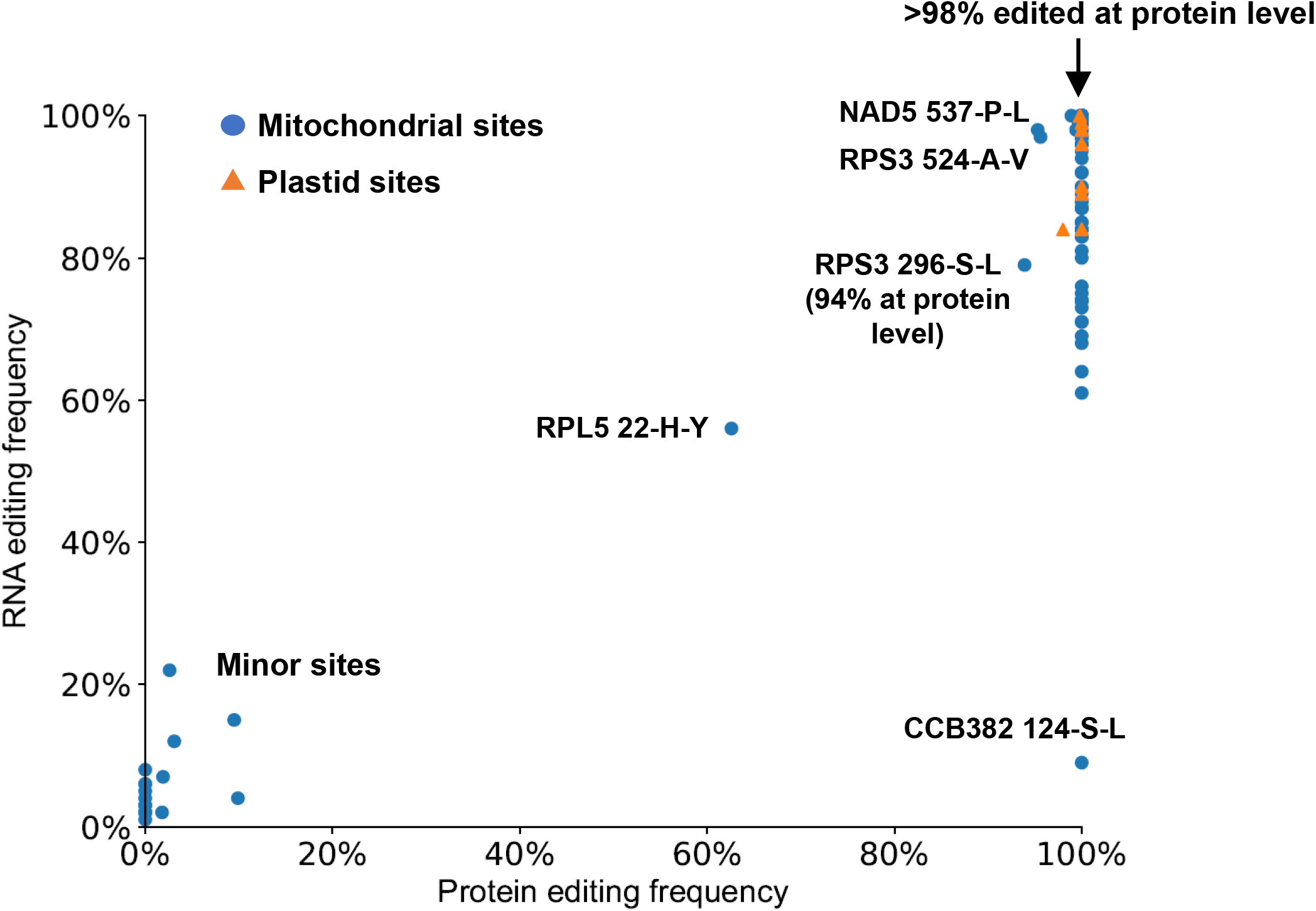
Quantitative relationship between editing at the RNA level (from RNAseq (Bentolila et al., 2013)) and the protein level determined in this study (only sites detected with at least 5 PSMs are shown). The underlying data are from Table 3 and 4.

Among the 36 mitochondrial sites whose editing extent is in the 50%-90% range as measured by RNA-seq (Bentolila et al., 2013) most (31 sites) are carried by transcripts encoding ribosomal proteins (Table 4 – sites in bold). Except for RPL5-22 and RPS3-296, all corresponding peptides carry only the edited version of the encoded amino acid (33 edited codons at a 100% occurrence). Given that ribosome profiling showed that most partially edited sites remain partially edited in polysome-associated mRNAs (Planchard et al., 2018), our results suggest that there must be some checkpoints either co-translationally or post-translationally that exclude the expression of partially edited transcripts. Sampling with a reduced number of PSMs could explain the absence of unedited translated codons for a small number of the sites; seven positions are covered by 1-61 PSMs (Table 4). However, the likelihood of not detecting unedited peptides for the remaining 28 positions is very low given the number of PSMs (107-2837). This observation is consistent with the prior report of the absence of proteins corresponding to *atp6* transcripts in petunia mitochondria (Lu and Hanson, 1994).

The only site in our study showing a significant amount of partially editing extent both at the transcript level and at the protein level (56% of edited transcripts and 63% of edited proteins, Table 4) is found in RPL5 at position 22. RPS12 is the only other plant mitochondrial protein previously described in the literature that is found both in edited and unedited translated forms (Lu et al., 1996; Phreaner et al., 1996). In maize, immunological analysis with two antibodies specific for the unedited or edited version of a peptide covering 3 partially edited sites at position 90, 95, and 97 demonstrated that both edited and unedited *rps12* translation products are present in mitochondria (Phreaner et al., 1996). However, only edited translation products accumulate in mature maize mitochondrial ribosomes. The authors speculated that the unedited RPS12 protein might have taken a separate function from its role in translation as a component of the ribosome. As a rationale for that hypothesis, RPS12 from *E. coli* is known to have nonspecific RNA binding protein activity and to facilitate intron splicing and RNA folding *in vitro* (Coetzee et al., 1994). In petunia, the same antibodies allow discrimination between the edited and unedited forms that result from the translation of two conserved editing sites with maize at position 90 and 95. Unlike maize, RPSI2 proteins recognized by both edited and unedited specific antibodies are present in the petunia mitochondrial ribosomal fraction (Lu et al., 1996). The authors also showed that unedited *rps12* translation products were detected in plant species other than petunia (*Cucumis pepo*, *Triticum aestivum*), demonstrating that the polymorphism in mitochondrial *rps12* expression is widespread. The extent of editing in Arabidopsis measured by RNA-seq was 88% for both rps12-90 and rps12-95 sites, and the MSMS analysis did only detect edited peptides (Table 4).

In *rpl5*, the partially edited tyrosine codon located at position 22 in Arabidopsis(position 27 in the alignment shown in Figure 6) is highly conserved. The tyrosine is genomically encoded in bacteria, green algae (*N. olivacea*), non-vascular plants comprising liverworts (*M. palacea*, *C. arguta*, *A. pinguis*) and mosses (*A. rugelii*, *C. dendroides*, *P. patens*, *P. commune*) and in some class of vascular plants, the lycopods or clubmosses represented by *Isoetes engelmannii*. In angiosperms (*O. sativa*, *S. latifolia*, and *A. thaliana*) editing of the first nucleotide in the codon for histidine (CAC) results in the creation of the codon encoding tyrosine (UAC). In the fern *Ophioglossum californicum* histidine is encoded in the mitochondrial *rpl5* gene as in the angiosperms, but evidence for editing at that position has not been established. Given the high level of conservation for tyrosine across different lineages, we can assume that its presence is required for a proper function of RPL5 in the ribosome. Experiments in bacteria have shown that RPL5 is a 5SrRNA-binding protein that is important for the incorporation of this ribosomal RNA into the 50S ribosomal subunit *in vitro* (Rohl and Nierhaus, 1982). In addition, the *E. coli rplE* gene encoding RPL5 is an essential gene as knockout of L5 gene is lethal (Korepanov et al., 2007). RPL5 has been shown to play a crucial role in the formation of the central protuberance containing 5S rRNA and proteins L5, L16, L18, L25, L27, L31, L33 And L35 during assembly of the large ribosomal subunit in the bacterial cell (Korepanov et al., 2012). These observations indicate the importance of this ribosomal protein for the ribosome formation and functioning. The existence of protein variants of RPL5 in Arabidopsis generated by editing at a position that is phylogenetically conserved constitutes a puzzling result. However, the binding of RPL5 to the 5S rRNA is mediated through one of the loops of this small RNA (Wang et al., 2013). It has been shown *in vitro* that RPL5 from *Arabidopsis* is able to bind specifically to the potato spindle tuber viroid RNA that possesses a similar loop as the 5S rRNA (Eiras et al., 2011). One can then speculate that like for RPS12, RPL5 might have taken on other functions in the mitochondrion than solely its involvement in the translation apparatus. Resolving the function of this protein polymorphism will require finding out whether both protein forms, histidine/unedited and tyrosine/edited are incorporated in the mitochondrial ribosome.

**Figure 6.**
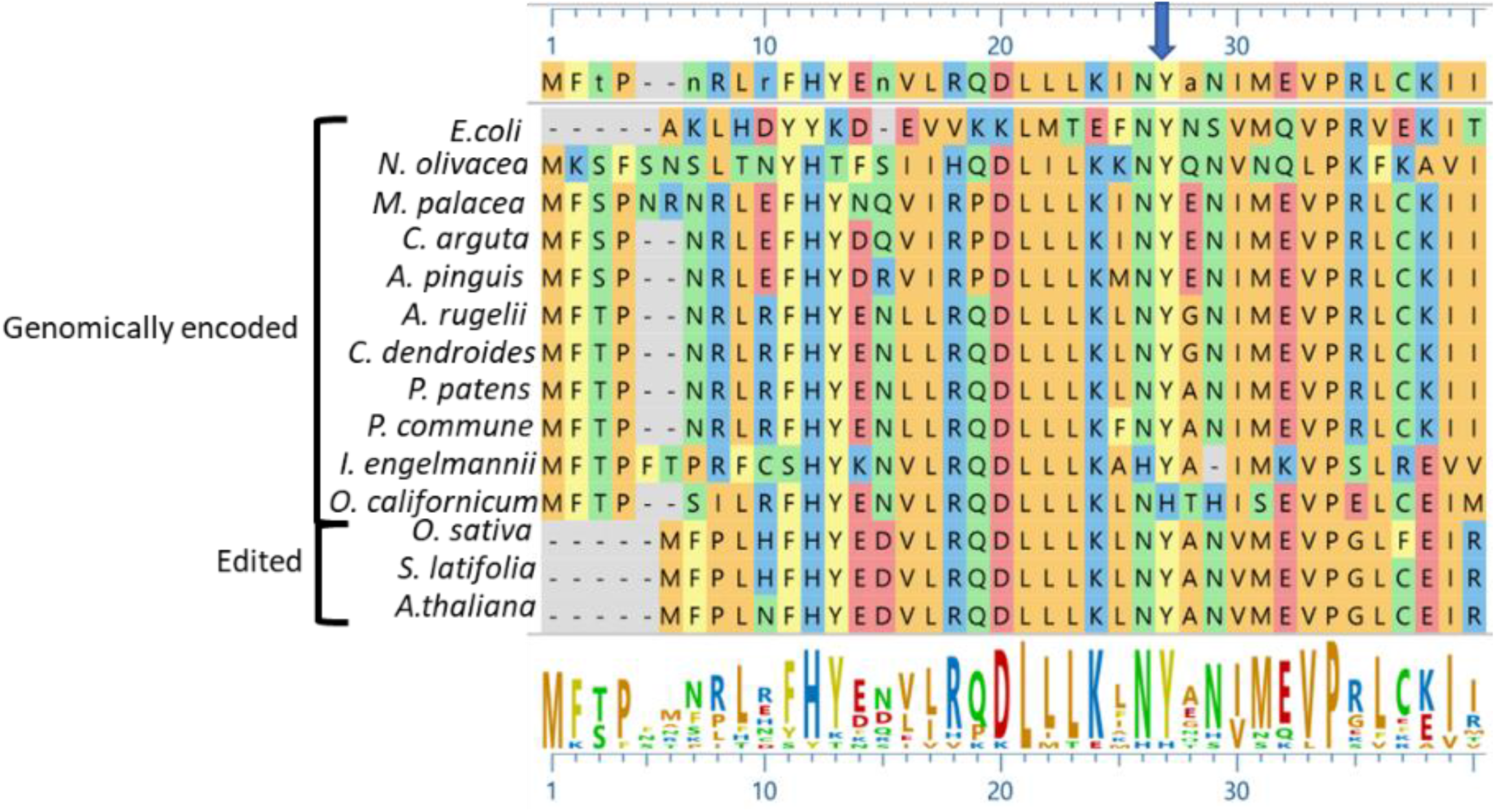
Alignment of the N terminus of RPL5 proteins in a wide range of species (bacteria, mammals, fungi, mosses, liverworts, ferns, kiwi, monocotyledons (*O. sativa*) and dicotyledons (*S. latifolia*, *A. thaliana*)). With the exception for the fern *O. californicum,* the conserved tyrosine that is either directly encoded or generated by RNA editing is indicated by an arrow.

In this study, we have observed that most of proteins found in the Arabidopsis organelles are translated from edited transcripts suggesting some control either co-translationally or post–translationally that excludes the presence of unedited proteins. In that respect, our results support the *raison d’être* of editing as a corrective mechanism which allows production of functional proteins. Nevertheless, we have also uncovered one instance of a protein polymorphism generated by editing in an essential protein, RPL5. Whether this polymorphism has an impact on the function(s) of RPL5 remains to be investigated.

### PTMs of the plastid– and mitochondrial encoded proteomes

Plant proteins undergo many PTMs to regulate protein function and stability (Friso and van Wijk, 2015; Grabsztunowicz et al., 2017) Here we reported on four of the most important and studied PTMs, *i.e.* N-terminal protein acetylation, lysine (side chain) acetylation, phosphorylation (on S,T,Y) and ubiquitination, based on the large scale reprocessing and searching of MSMS data. Importantly, with the exception of one observation for ubiquitination (plastid RPS7-K13), we did not identify any ubiquitination for plastid– or mitochondrial encoded proteins (but we identified ∼1000 ubiquitinated nuclear-encoded proteins). This is in line with our recent re-assessment (van Wijk et al., 2023) of a recent paper that claimed extensive polyubiquitination in the chloroplasts (Sun et al., 2022). The lack of ubiquitination of plastid-encoded proteins is consistent with the lack of known E1-E2-E3 enzymes involved in ubiquitin activation and ligation. We also did not observe any N-terminal acetylation of mitochondrial-encoded proteins consistent with a previous report (Huang et al., 2009) and the lack of known acetyl aminotransferases in mitochondria (Pozoga et al., 2022). In contrast, 17 of the plastid-encoded proteins were N-acetylated at amino acid position 1 and 2, consistent with the presence of multiple acetyl amino transferases (Pozoga et al., 2022) and previous results. However, the functional significance of these N-terminal acetylations is not yet demonstrated, but is likely to impact protein stability (*e.g.* through an N-degron pathway (Bouchnak and van Wijk, 2019; Aguilar Lucero et al., 2021) or impact protein-protein interactions. The lack of observed N-terminal acetylation for the mitochondrial-encoded proteins, despite searching such a large set of MSMS data (∼∼257 million), further supports the notion that this PTM does not occur for mitochondrial-encoded proteins, and it also indicates a very low false discovery rate.

Lysine ε-amine acetylation is driven by the presence of acetyl molecules as part of fatty acid metabolism, and likely contributes to fine-tuning the activity of central metabolic enzymes, respiration and photosynthetic activity with likely impact on plant acclimation capacity (Koskela et al., 2018; Balparda et al., 2022; Fussl et al., 2022). Here we identified extensive lysine acetylation of 16 plastid-encoded proteins, and two mitochondrial proteins (ATP1 and NAD1) (4 sites). MSMS analysis of leaf proteome for Arabidopsis, and other species (*e.g.* soybean rice) suggests that thousands of plant proteins undergo lysine acetylation (Bienvenut et al., 2020; Li et al., 2021). Our data show that also plastid-encoded proteins are targets of protein acetylases. It remains to be determined what the biological impact is on these organellar proteins and which GNATs are responsible for these PTMs.

Phosphorylation has been observed for both plastid and mitochondrial localized proteins, in particular for nuclear-encoded proteins imported into these organelles (van Wijk et al., 2014; Willems et al., 2019; Zhang et al., 2021) and the thylakoid membrane system is well known to be regulated by phosphorylation (Rantala et al., 2020; Longoni and Goldschmidt-Clermont, 2021) The single phosphorylated mitochondrial protein (RPS3 (pS112) is important for mitochondrial biogenesis as loss of RPS3 splicing results in strong phenotypes (Sakamoto et al., 1996), but a role of phosphorylation of RPS3 has yet to be determined. The 12 plastid-encoded phosphorylated proteins are part of Photosystem II, Photosystem I, the ATP-synthase complex, the 30S ribosome, RBCL and TIC214. Thylakoid protein phosphorylation is governed by two state transition protein kinases (STN7,8), two protein phosphatases (PPH1/TAP38 and PBCP), whereas plastid casein kinase (pCKII) is involved in the broader network of plastid protein phosphorylation (Rantala et al., 2020; Longoni and Goldschmidt-Clermont, 2021). RBCL (pT333) and RPS7 (pS93) phosphorylation sites were also detected and shown to be (mostly) dependent of kinase STN7 Schonberg, 2017 #20919}. Consistently, whereas we observed multiple phosphosites in RBCL, pT333 was by far the most frequently observed (68 PSMs). Of the previously reported N-terminal threonine p-sites on the plastid-encoded PSII core proteins, D1, D2, CP43 and PSBH upon exposure to high light (Rantala et al., 2020; Longoni and Goldschmidt-Clermont, 2021), we did identify two closely spaced phosphorylated threonines on PSBH (T3 and T5) (Supplemental Data Set 1). We also identified phosphorylation on four of the plastid-encoded subunit of the plastid ATP-synthase complex (CF1A,B and CFO-I,III) of which the two CF1 subunits were also previously found to be phosphorylated (Sattari Vayghan et al., 2022).

## CONCLUSIONS

This study is an important milestone in defining the plastid– and mitochondrial-encoded proteins with correct protein sequences, RNA editing state and coverage of several physiological PTMs in Arabidopsis Col-0. The survey of organelle editing at a proteome wide level shows that editing is nearly always required for stable protein accumulation, even for sites where RNA editing well below 100%. These findings support the maintenance of this complex phenomenon in plants solely for the purpose of correcting deleterious mutations at the RNA level. This information should be incorporated in the next Arabidopsis genome build and annotation and it will simplify MS-based proteome analysis of these organelle-encoded proteins. This study also demonstrates the benefits of reanalyzing publicly available MSMS raw data sets at a large scale and shows the importance of careful verification of the underlying MS data for low frequency events.

## MATERIALS AND METHODS

### The Arabidopsis PeptideAtlas build #2

The analysis in this study was based upon the ensemble results of the 2022-06 build of the Arabidopsis PeptideAtlas, as described in detail in (van Wijk et al., 2023). Briefly, the raw data files from 115 datasets were downloaded from ProteomeXchange data repositories (Deutsch et al., 2023) and reprocessed with the Trans-Proteomic Pipeline (Deutsch et al., 2023) using parameters appropriate to each dataset. This resulted in the identification of over 70 million spectra, nearly 0.6 million peptides, and 18267 proteins (out of 27559) at the highest confidence level.

### Creating the protein search space and search strategy, allowing for partial editing

The protein search space included all protein sequences in Araport11, TAIR10, UniProtKB, Refseq and others, as described in (van Wijk et al., 2023). We also included the consensus set of 79 plastid-encoded and 33 mitochondrial-encoded non-redundant protein sequences (Tables 1 and 2) to address mistakes and redundancies in plastid– and mitochondrial ATGC and ATMG identifiers in Araport11 (same as in TAIR10) (as explained further in RESULTS). We also included variants of protein sequences for those plastid– and mitochondrial encoded proteins that are predicted to be affected by RNA editing. For the mitochondrial-encoded proteins, we included 420 editing sites in 29 mitochondrial-encoded proteins and two ORFs, most of which are described in (Sloan et al., 2018) whereas we included edited sequences for 17 plastid-encoded proteins that included 31 amino acid changes and generation of one start methionine. These organellar-encoded sequences included unedited sequences, completely edited sequences. If predicted editing sites were sufficiently close together to appear in a single peptide (*i.e.* within ∼30 amino acids of each other), we also included additional sequences that contained all possible permutations of non-edits and edits, to be able to detect instances of partial editing in one proteoform. This resulted in the addition of 10,368 sequences for plastid– and mitochondrial encoded variants to the search database. These non-synonymous editing sites were carefully assembled based on information from publications, public sequence databases and discussions with experts in plant organelle RNA editing, in particular TAIR (Araport11 and TAIR10), UniProtKB, NCBI-Refgen, *A. thaliana* Col-0 ecotype mitochondrial data set from (Sloan et al., 2018), as well as plant RNA editing databases PREPACT3.0 (http://www.prepact.de/prepact-main.php), REDIdb RNA editing database (http://srv00.recas.ba.infn.it/redidb/) and the Plant editosome database (https://ngdc.cncb.ac.cn/ped/) (Lenz et al., 2018) (Lo Giudice et al., 2018) (Li et al., 2019).

### Manual verification of editing sites detected at low frequency

To ensure that editing sites observed in the proteome at low frequency (in their unedited or edited form) were correctly identified, matched MS/MS spectra were manually evaluated in PeptideAtlas build #2 to verify for presence of y and b ions that specifically identified residue positions that could be affected by editing. For this editing site analysis, we also restricted possible variable mass modifications to methionine oxidation (but included semi-tryptic peptides) to reduce possible false positive editing sites. The final reported results in this publication are made after these evaluations; hence, the spectra that were identified by the software but did not pass manual validation are still available in PeptideAtlas.

## ACKNOWLEDGEMENTS AND FUNDING

We thank the plant organelle research community, in particular Ian Small, Josh Hazelwood, Etienne Meyer, Ralph Bock for information and feedback on organellar genes and editing sites. This research was support by a grant from NSF-IOS 1922871 to KJVW and EWD.

## AUTHOR CONTRIBUTIONS

TL and ZS carried out the MS searches and PeptideAtlas data loading, supervised by EWD, and assembled the search results. ML contributed to data analysis. LM and ZS developed the PeptideAtlas web interface. QS helped assemble the protein search space. EWD, KJVW and SB developed the project, evaluated outcomes, and wrote the paper.

## SUPPLEMENTAL DATA

**Supplemental Data Set S1**. Detection of four physiological important PTMs, N-terminal protein α-amine acetylation, lysine ε-amine acetylation, phosphorylation and ubiquitination.

